# Agrochemical control of gene expression using evolved split RNA polymerase. II

**DOI:** 10.1101/2024.04.02.587689

**Authors:** Yuan Yuan, Jin Miao

**Affiliations:** Department of Neurophysiology and Neuropharmacology, Institute of Special Environmental Medicine and Co-innovation Center of Neuroregeneration, Nantong University, Nantong, 226019, China; Duke Kunshan University, 8 Duke Avenue, Kunshan, Jiangsu Province, 215316, China

**Author notes:** Corresponding Author: Jin Miao, Duke Kunshan University, 8 Duke Avenue, Kunshan, Jiangsu Province, 215316, China.

## Abstract

Agrochemical inducible gene expression system provides cost-effective and orthogonal control of energy and information flow in bacterial cells. However, the previous version of Mandipropamid inducible gene expression system (Mandi-T7) became constitutively active at room temperature. We moved the split site of the eRNAP from position LYS179 to position ILE109. This new eRNAP showed proximity dependence at 23°C, but not at 37°C. We built Mandi-T7-v2 system based on the new eRNAP and it worked in both *Escherichia coli and Agrobacteria tumefaciens*. The modified eRNAP when combined with the leucine zipper-based dimerization system, behaved as a cold inducible gene expression system. Our new system provides a means to broaden the application of agrochemicals for both research and agricultural application.

## Introduction

Agrochemicals are key to health promotion and growth management in modern agriculture. The development of biosensors for agrochemicals opens new avenues to remote control of cell behavior (Park, 2015, Zimran, 2022, Park, 2024). One of the possible applications is to use agrochemicals to control gene expression of bacteria associated with plants or animals. Mandipropamid is an oomycete fungicide. We recently reported the development of Mandipropamid inducible gene expression system (Mandi-T7) using a protein proximity detection platform based on evolved split T7 RNA polymerase (eRNAP). Repurposed plant Abscisic acid receptor proteins will sense agrochemical Mandipropamid and will be brought into proximity upon Mandipropamid binding (Park, 2015). This in turn activates the eRNAP and leads to expression of genes driven by the T7 promoter (Figure 1A). Wild type T7 RNAP split at site 179 is proximity-independent and was used to construct transcriptional logic gates (Shis, 2013 and Segall-Shapiro, 2014). The simultaneous assembly of the split T7 RNAP was abolished through directed evolution (eRNAP) and the eRNAP became proximity-dependent (Pu, 2017). Several small molecule inducible expression systems were developed based on this eRNAP platform, such as Abscisic acid and Rapamycin (Pu, 2018), Mandipropamid (Yuan, 2022), and Rapalog (Martin, 2023). The molecular variety of this eRNAP biosensor platform has been further expanded by fusion with the variable domains of antibodies (Komatsu, 2023) or with cell-pole organizing proteins to achieve asymmetric gene expression (Lin, 2021).

**Figure 1.**
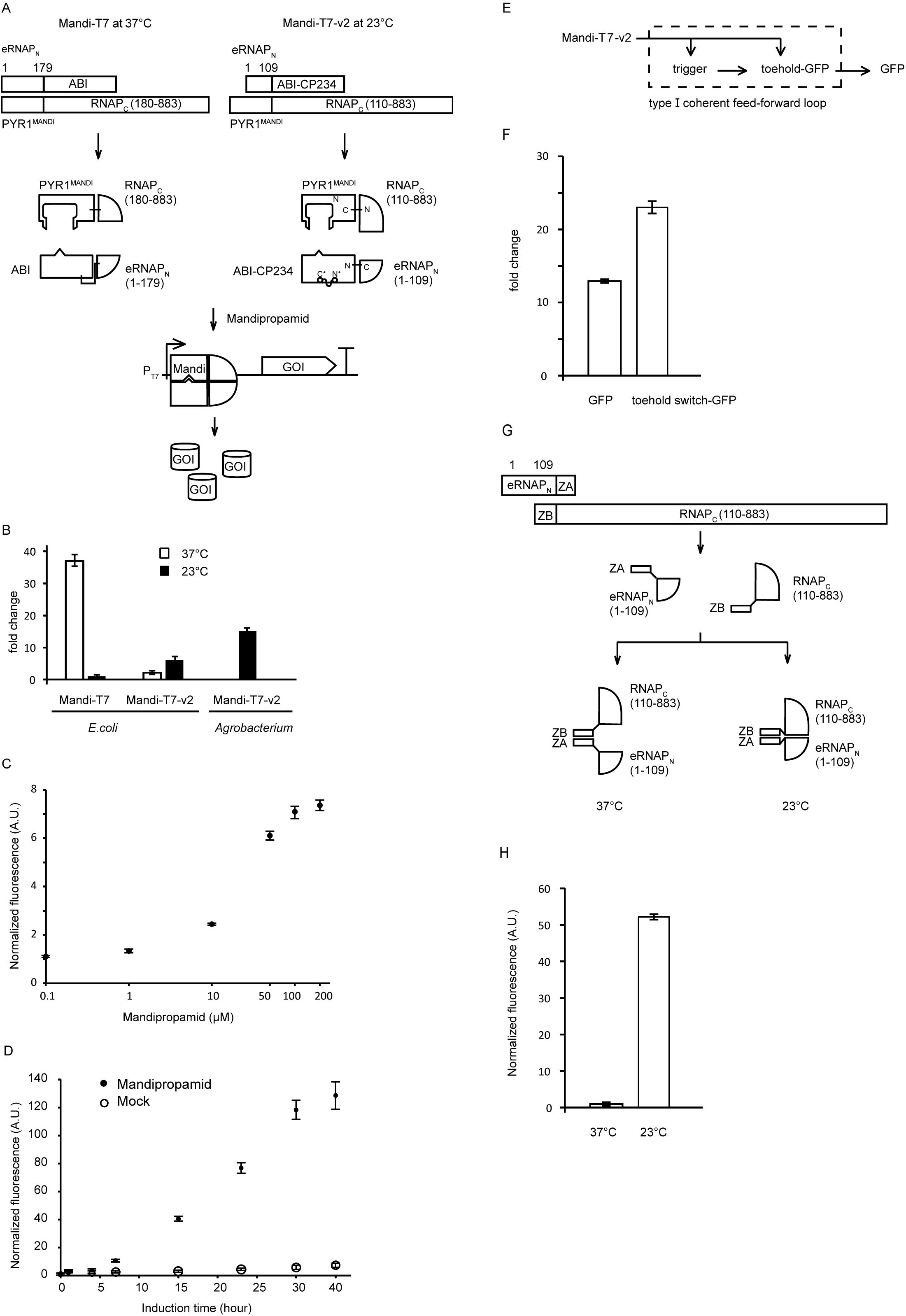
Engineering the Mandi-T7-v2 system. (A) Comparison of the Mandi-T7 and Mandi-T7-v2 systems. The length of different parts of fusion proteins were drawn in scale. Bent arrow: T7 promoter; large T-shape: termitor. (B) fold change of reporter gene expression after induction at 23 □or 37 □. (C) Dose response analysis of Mandi-T7-v2 in E.coli after induction at 23 □for 6 hours. (D) kinetic analysis of Mandi-T7-v2 system at 23 □. (E) schematic of toehold switch incorporated into Mandi-T7-v2 induction system. (F) fold change of report gene expression. (G) schematic of leucine zipper peptides ZA/ZB mediated dimerization of eRNAP2. (H) temperature dependent expression of the reporter gene.

The Mandi-T7 system was initially tested at 37°C.The failure of Mandi-T7 to work at room temperature hampers engineering initiatives in other bacteria. Here we report alleviation of this issue by adopting a new split site of T7 eRNAP. The new system Mandi-T7-v2 works at 23°C in both *E*.*coli* and *Agrobacterium*. The modified eRNAP is also compatible with a leucine zipper peptide-based dimerization system and may be used with other chemical induced dimerization systems as well.

## Material and Methods

### Plasmid construction

The eRNAP_N_ (derived from 1-109 aa of T7 RNAP), the T7 RNAP_C_ (110-883 aa of T7 RNAP), and ZA/ZB fragments were synthesized and cloned into the plasmid pJM1B6 (Yuan, 2022) by GenScript Biotechnology (Nanjing, China). The pVS1 origin from pCAMBIA1301 (GenBank: AF234297.1, 2488 bp - 6266 bp), the Mandi-T7-v2 driver cassette, and the T7p::mcherry effector cassette were assembled into the pGM1190 plasmid (Addgene #69994) backbone to enable single-plasmid expression in *Agrobacterium*. Detailed information of these genetic parts and plasmids was listed in Table S1 and S2.

### Mandipropamid responsive assay for evolved split T7 RNA polymerase

The response to Mandipropamid was assayed as previously reported. Briefly, the strain Top10 (TIANGEN biotech., Beijing, China) was transformed with the driver and reporter plasmids. Single colonies were inoculated into SOC medium supplemented with Ampicillin (100mg/mL) and Spectinomycin (50 mg/mL), and the mixture was allowed to grow at 37 °C. The overnight culture was transferred to fresh medium with antibiotics at 1:400 ratio and incubated for 3 hours at 37 °C. Then induction by Mandipropamid was tested at both 23°C and 37 °C. Mandipropamid (sc-235565, Santa Cruz) was added as the inducer, and DMSO as the solvent control. After incubating for 3 hours at 37 °C or 6 hours at 23°C, 100 µL of each sample was transferred to a 96-well plate. The florescence signal (GFP, Ex: 488, Em: 510; mcherry, Ex: 587, Em: 630) and OD_600_ of the culture were then measured using a ThermoFisher Varioskan LUX plate Reader. For *Agrobacterium tumefaciens*, the strain LBA4404 (Weidibio, Shanghai, China) was transformed with the plasmid pJM-Mandi-T7-Ag and grown at 28°C. Single colonies were inoculated SOC medium supplemented with Apramycin (50 mg/mL) and grown at 28°C. The overnight culture was transferred to fresh medium with antibiotics at 1:100 ratio and incubated for 3 hours at 28 °C. Induction was performed overnight at 23°C after adding Mandipropamid or DMSO.

### Cold responsive assay of split eRNAP fused with leucine zipper peptides

The *E. coli* strain Top10 was used. Single colonies were inoculated SOC medium supplemented with Ampicillin (100mg/mL) and Spectinomycin (50 mg/mL) and grown at 37 °C. The overnight culture was transferred to fresh medium with antibiotics at 1:400 ratio and incubated for 3 hours at 37 °C. Following this, half the culture was incubated at 23°C, the other half stayed at 37 °C. After 3hr at 37 °C or overnight at 23°C, 100 µL of each sample was transferred to a 96-well plate. The florescence signal (GFP, Ex: 488, Em: 510; mcherry, Ex: 587, Em: 630) and OD_600_ of the culture were measured using a ThermoFisher Varioskan LUX plate Reader.

### Time course fluorescence measurement

*E. coli* Top10 strain containing Mandi-T7_v2 and reporter plasmids was generated as mentioned above.

Single colonies were used to inoculate SOC medium supplemented with Ampicillin (100 mg/mL) and Spectinomycin (50 mg/mL) and grown at 37 °C overnight. The Overnight culture was transferred to fresh medium with antibiotics at 1:100 ratio. Afterwards, either Mandipropamid (50 mM, sc-235565, Santa Cruz) or DMSO (solvent control) was added at a 1:1000 ratio. The culture was incubated at 23°C for 40 hours. Samples were harvested after 1 hour, 4 hours, 7 hours, 15 hours, 23 hours, 30 hours and 40 hours. The florescence (GFP: Ex: 488, Em: 510; mcherry: Ex: 587, Em: 630) and OD_600_ of 100 µL of each sample were measured using a ThermoFisher Varioskan LUX plate Reader.

## Results

To apply Mandi-T7 to bacteria living at room temperature, we characterized Mandi-T7 at 23°C . The results showed that fluorescence signal of the report gene could be detected regardless of the presence of the inducer (Figure 1B and Supplemental Figure 1), indicating that the eRNAP relapsed into self-assembly at 23°C. Restoration eRNAP’s proximity dependency at 23°C will enable Mandi-T7 to work at 23°C. We speculated that a new split site in the N terminal region of T7 RNA polymerase, not previously reported by the bisection mapping efforts (Segall-Shapiro, 2014), might prevent self-assembly at 23°C. We scrutinized the structure of the T7 RNAP N terminal region and selected ILE109 as our candidate, because ILE109 is exposed at the surface and splitting at ILE109 / LYS110 will break the connection between the Arginine loop, which is essential for binding upstream AT-rich region of the T7 promoter, with the rest of the T7 RNA polymerase (Supplemental Figure 2).

To assess the potential improvement in induction at 23°C, we modified the driver module of the Mandi-T7 system (Figure 1A). The split site was changed to ILE109/LYS110 while retaining the point mutations in the N-terminal region of the eRNAP (F21L, L32S, E35G, R57C, E63K, K98R, Q104K, Q107K) as leverage for flexibility. Induction for 6 hours at 23°C yielded modest fold change over the control (Figure 1B). Unexpectedly, induction by Mandipropamid at 37°C was abolished by adopting the new split site (Figure 1B). Encouraged by these findings, we named this modified eRNAP as eRNAP2, and our new Mandipropamid inducible expression system as Mandi-T7-v2. We proceeded to characterize the induction with different concentrations of Mandipropamid. We observed maximum induction by 200 µM Mandipropamid at 23°C (Figure 1C). We further evaluated the kinetic characteristics of the Mandi-T7-v2 at 23°C over 30 hours. We observed induction signal after 1 hour and continuous increase over 30 hours (Figure 1D and Supplemental Figure 1). However, the fluorescence signal of the non-induction samples accumulated at the same time, resulting in marginal increase in fold induction after 23 hours (Figure 1D).

To increase the dynamic range without further engineering the Mandi-T7-v2 system, we tried to incorporate the toehold switch into the reporter module (Figure 1E), which have been shown to improve the performance of T7 based inducible expression systems through a coherent type 1 feed-forward loop (Hwang, 2021; Greco, 2021). Indeed, incorporation of the toehold switch could increase the dynamic range by two-fold (Figure 1F).

To test whether Mandi-T7-v2 can be applied to other bacteria, we tried Mandi-T7-v2 in *Agrobacterium tumefaciens*, a plant pathogen and medium of T-DNA transformation. To generate a single-plasmid based induction system, we assembled the Mandi-T7-v2 driver module, the reporter module and the pVS1 replication origin together. We tested the Mandipropamid induction in *Agrobecterium* at 23°C.

The result showed 15-fold induction (Figure 1B and Supplemental Figure 1), suggesting that the Mandi-T7-v2 system works in other bacteria as well.

To test whether induction at 23°C but not at 37°C is an inherent property of the new eRNAP2 or depends on the fusion to the ABI-PYR1^MANDI^ protein pair. We replaced the ABI-PYR1^MANDI^ pair with leucine zipper peptides ZA and ZB (Figure 1G), one of the model dimerization systems which will lead to spontaneous dimerization and has been used for developing the evolved RNAP system before (Pu, 2017; Pu, 2018). We tested the ZA/ZB-eRNAP2 system at both 23°C and 37°C. The results showed 50-fold higher expression at 23°C over 37°C (Figure 1H and Supplemental Figure 1), reminiscent of a cold inducible expression system. This result indicates the restored proximity dependency of the eRNAP2 at 23°C does not require specific fusion partners.

## Discussions

We are fortunate to have found a new split site of eRNAP, which enables chemical inducible activity control at 23°C. Induction around 23°C will be an advantage for controlling gene expression in bacteria living around room temperature. This new eRNAP2 system can also be adapted to other proximity-dependent systems.

When we tested the performance of Mandi-T7 system at room temperatures, we found constitutive activity without inducibility, indicating the disrupted spontaneous assembly of eRNAP at 37°C was restored at room temperature. The temperature sensitivity of the eRNAP platform may be explained by its origin in the directed evolution system running at 37°C (Pu, 2017).

Several split sites of T7 RNA polymerase have been identified by the bisection mapping, like around 67, 301, 563, and 763 (Segall-Shapiro, 2014). Except split sites 563, which has also been used for engineering the thermo-repressible split T7 RNAP (Chee, 2022), and the blue light-inducible systems (Han, 2017 and Dionisi, 2022), these split sites have not been developed into general or specific biosensor platforms. Our work suggests that it is worth exploring new split sites of T7 RNA polymerase for specific applications.

We incorporated toehold switch into our simple gene circuit to improve dynamic range. Further directed-evolution efforts may be required to reduce the background activity of the Mandi-T7-v2 system.

The ABI-PYR1 pair has been recently expanded to be sensors of other agrochemicals (Zimran, 2022; Park, 2024). Our work will readily help develop new inducible gene expression systems of these agrochemicals.

## Acknowledgement

We would like to thank Professor Linfeng Huang (Duke Kunshan University) for his discussions and generous sharing equipment and reagents. We also want to express our gratitude to Dr. Jinyue Pu and Professor Bryan Dickinson of the University of Chicago for their sharing plasmid information of the original eRNAP system.

---

**Supplemental figure 1.** cell pellet of E.coli and Agrobacterium culture post induction.

Samples were placed on the sill of a window with normal day light. +: Mandipropamid (50 μM final); -: DMSO.

**Supplemental figure 2.** the ILE109 split site of T7 RNA polymerase.

The N terminal region (1-109) of the T7 RNA polymerase was highlighted in dark purple. The Arginine loop responsible for binding the AT-rich region was encircled by a red square. The cartoon was generated from the structure of T7 RNA polymerase – T7 promoter complex (PDB ID: ICEZ).

