## Supplementary figures and images for "Agrochemical control of gene expression using evolved split RNA polymerase. II"

### supplemental figure 1

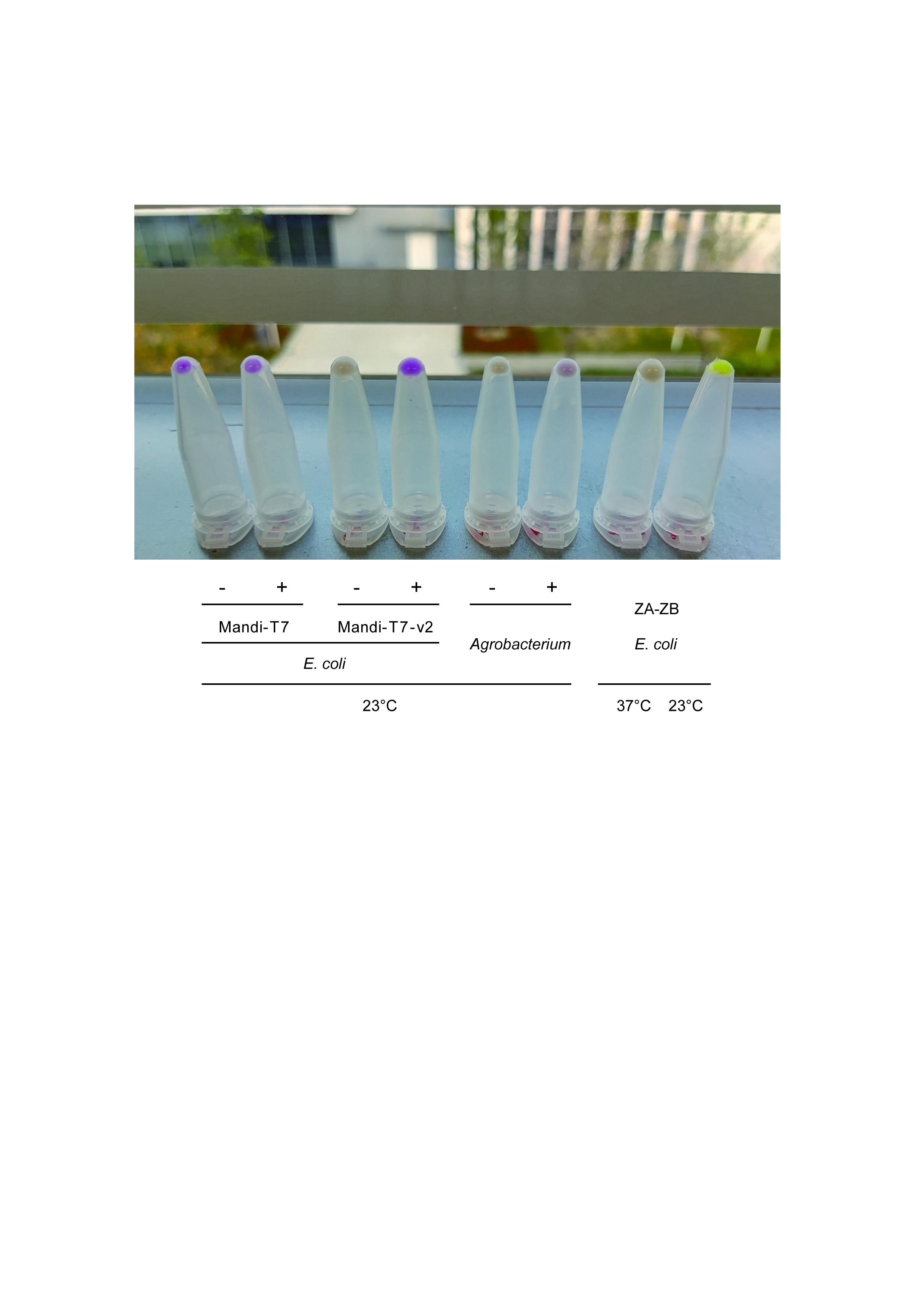

### supplemental figure 2

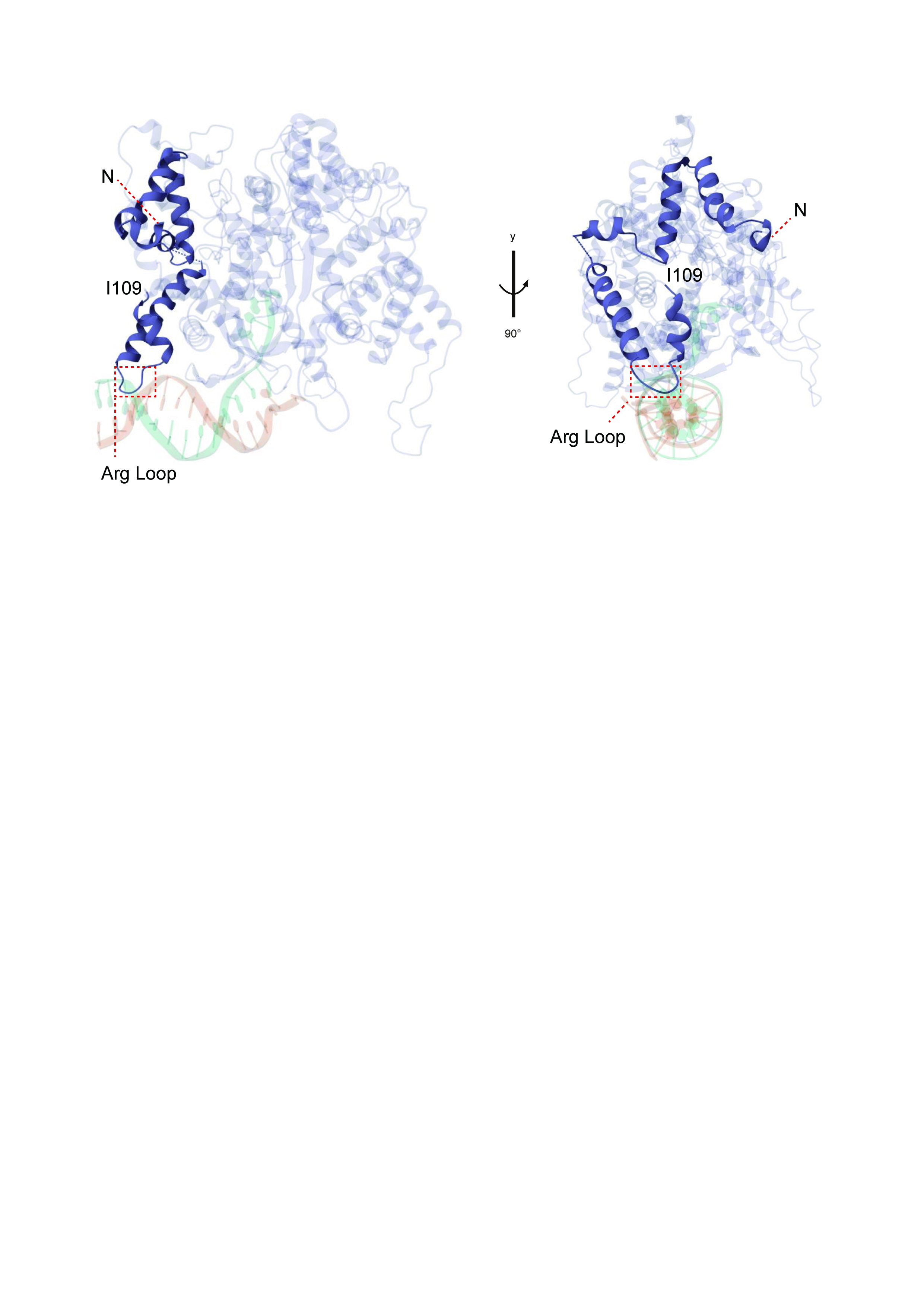
