## supplemental table 1 for "Agrochemical control of gene expression using evolved split RNA polymerase. II"

Table S1: sequence and source of genetic parts

| Name | Sequence | Source |
| --- | --- | --- |
| T7-eRNAP_N_ | MNTINIAKNDFSDIELAAIPLNTLADHYGERSARGQLALEHESYEMGEARFRKMFECQLKAGKVADNAAAKPLITTLLPKMIARINDWFEEVKAKRGRRPTAFKFLKEI  (from d5-19: F21L, L32S, E35G, R57C, E63K, K98R, Q104K, Q107K) | Pu, 2018 |
| T7-RNAP_C_  (110-883) | KPEAVAYITIKTTLACLTSADNTTVQAVASAIGRAIEDEARFGRIRDLEAKHFKKNVEEQLNKRVGHVYKKAFMQVVEADMLSKGLLGGEAWSSWHKEDSIHVGVRCIEMLIESTGMVSLHRQNAGVVGQDSETIELAPEYAEAIATRAGALAGISPMFQPCVVPPKPWTGITGGGYWANGRRPLALVRTHSKKALMRYEDVYMPEVYKAINIAQNTAWKINKKVLAVANVITKWKHCPVEDIPAIEREELPMKPEDIDMNPEALTAWKRAAAAVYRKDKARKSRRISLEFMLEQANKFANHKAIWFPYNMDWRGRVYAVSMFNPQGNDMTKGLLTLAKGKPIGKEGYYWLKIHGANCAGVDKVPFPERIKFIEENHENIMACAKSPLENTWWAEQDSPFCFLAFCFEYAGVQHHGLSYNCSLPLAFDGSCSGIQHFSAMLRDEVGGRAVNLLPSETVQDIYGIVAKKVNEILQADAINGTDNEVVTVTDENTGEISEKVKLGTKALAGQWLAYGVTRSVTKRSVMTLAYGSKEFGFRQQVLEDTIQPAIDSGKGLMFTQPNQAAGYMAKLIWESVSVTVVAAVEAMNWLKSAAKLLAAEVKDKKTGEILRKRCAVHWVTPDGFPVWQEYKKPIQTRLNLMFLGQFRLQPTINTNKDSEIDAHKQESGIAPNFVHSQDGSHLRKTVVWAHEKYGIESFALIHDSFGTIPADAANLFKAVRETMVDTYESCDVLADFYDQFADQLHESQLDKMPALPAKGNLNLRDILESDFAFA* | pET28 |
| ZA | ALKKELQANKKEIAQLKWEIQALKKELAQ | Pu, 2018 |
| ZB | MASEQLEKKLQALEKKLAQLEWKNQALEKKLAQ | Pu, 2018 |
| Toehold Switch trigger | GGGCTCGATCACTAATCTGATCGAGACGAACATACCTACCTTCATTATCTTACTTGT | Green, 2017 |
| Toehold Switch | GGGAGTAAGATAATGAAGGTAGGTATGTTAAACTTTAGAACAGAGGAGATAAAGATGAACATACCTACGAACCTGGCGGCAGCGCAAAAG | Green, 2017 |

Table S2: schematics of vectors

| Name | schematic | Used in |
| --- | --- | --- |
| pJM-3A03 | 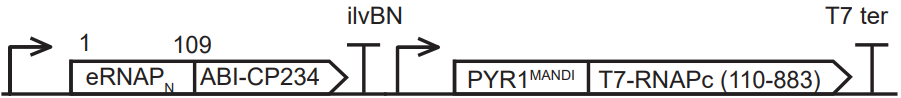 | Fig1B, 1C, 1D, 1F |
| pCDF-T7-mcherry | 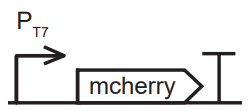 | Fig1B, 1C, 1D, 1H |
| pJM-Mandi-T7-Ag | 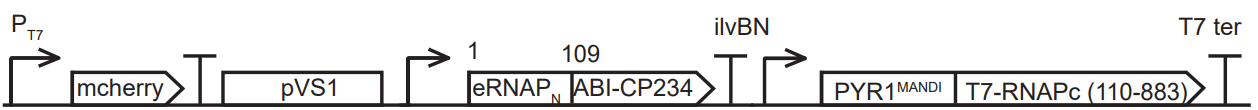 | Fig 1B |
| pCDF-Toehold | 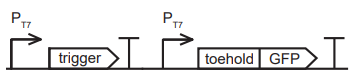 | Fig 1F |
| pJM-ZAZB | 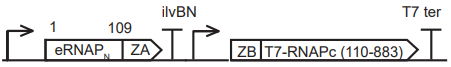 | Fig 1H |
