## supplemental table 2 for "Agrochemical control of gene expression using evolved split RNA polymerase. II"

Table S2. Schematic of plasmids

| Name of plasmids | Schematic | application |
| --- | --- | --- |
| Mandi-T7-v2 | 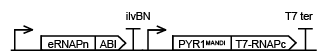 | Fig 1B, 1C, 1D, 1F |
| pCDF-T7-mcherry | 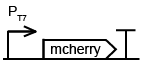 | Fig 1B, 1C, 1D |
| pJM-Mandi-T7-Ag | 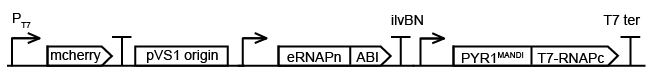 | Fig 1B |
| pCDF-T7-sfGFP | 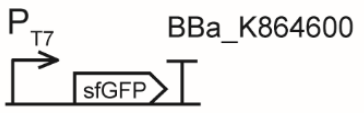 | Fig 1F |
| pCDF-T7-toehold-GFP | 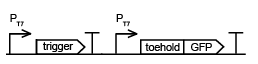 | Fig 1F, 1H |
| ZA-ZB-eRNAP2 | 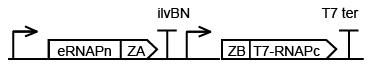 | Fig 1H |
